# Influence of human cytomegalovirus glycoprotein O polymorphism on the inhibitory effect of soluble forms of trimer- and pentamer-specific entry receptors

**DOI:** 10.1101/778241

**Authors:** Nadja Brait, Tanja Stögerer, Julia Kalser, Barbara Adler, Ines Kunz, Max Benesch, Barbara Kropff, Michael Mach, Elisabeth Puchhammer-Stöckl, Irene Görzer

**Affiliations:** Center for Virology, Medical University of Vienna, Vienna, Austria; Max von Pettenkofer-Institute for Virology, Ludwig-Maximilians-University Munich, Munich, Germany; Virologisches Institut, Klinische und Molekulare Virologie, Friedrich-Alexander Universität Erlangen-Nürnberg, Erlangen, Germany

## Abstract

Human cytomegalovirus (HCMV) envelope glycoprotein complexes, gH/gL/gO-trimer and gH/gL/UL128L-pentamer, are important for cell-free HCMV entry. While soluble Nrp2-Fc (sNrp2-Fc) interferes with epithelial/endothelial cell entry through UL128, soluble PDGFRα-Fc (sPDGFRα-Fc) interacts with gO thereby inhibiting infection of all cell types. Since gO is the most variable subunit we investigated the influence of gO polymorphism on the inhibitory capacities of sPDGFRα-Fc and sNRP2-Fc.

Accordingly, gO genotype 1c (GT1c) sequence was fully or partially replaced by gO GT2b, GT3, GT5 sequences in TB40-BAC4-luc background. All mutants were tested for fibroblast and epithelial cell infectivity, for virions’ gO and gH content, and for infection inhibition by sPDGFRα-Fc and sNrp2-Fc.

Full-length and partial gO GT swapping may strongly alter the virions’ gO and gH levels associated with enhanced epithelial cell infectivity. All gO GT mutants except recombinant gO GT1c/3 displayed a near-complete inhibition at 1.25 μg/ml sPDGFRα-Fc on epithelial cells (98% versus 91%) and all on fibroblasts (≥ 99%). While gO GT replacement did not influence sNrp2-Fc inhibition at 1.25 μg/ml on epithelial cells (96%-98%), it rendered mutants with low gO levels moderately accessible to fibroblasts inhibition (20%-40%). In contrast to the steep sPDGFRα-Fc inhibition curves (slope >1.0), sNrp2-Fc dose-response curves on epithelial cells displayed slopes of ~1.0 suggesting functional differences between these entry inhibitors.

Our findings suggest that targeting of gO-trimer rather than UL128-pentamer might be a promising target to inhibit infectivity independent of the cell type, gO polymorphism, and gO/gH content. However, intragenic gO recombination may lead to moderate resistence to sPDGFRα-Fc inhibition.

**Importance:** Human cytomegalovirus (HCMV) is known for its broad cell tropism as reflected by the different organs and tissues affected by HCMV infection. Hence, inhibition of HCMV entry into distinct cell types could be considered as a promising therapeutic option to limit cell-free HCMV infection. Soluble forms of cellular entry receptor PDGFRα rather than those of entry receptor neuropilin-2 inhibit infection of multiple cell types. sPDGFRα specifically interacts with gO of the trimeric gH/gL/gO envelope glycoprotein complex. HCMV strains may differ with respect to the virions’ amount of trimer and the highly polymorphic gO sequence. In this study, we show that gO polymorphism rather than gO levels may affect the inhibitory capacity of sPDGFRα. The finding that gO intragenic recombination may lead to moderate evasion from sPDGFRα inhibition is of major value to the development of potential anti-HCMV therapeutic compounds based on sPDGFRα.

## Introduction

Human cytomegalovirus (HCMV) is a widely spread pathogen which may cause substantial harm in congenitally infected newborns and in patients undergoing severe immunosuppressive therapy (1). Natural HCMV transmission follows mainly through body fluids such as urine or saliva (2). Upon infection, HCMV is spread throughout the body infecting many of the major somatic cell types like fibroblasts, smooth muscle cells, endothelial cells, epithelial cells, neurons, and leukocytes (3).

Two virion envelope glycoprotein (gp) complexes of human cytomegalovirus, the trimer gH/gL/gO and the pentamer gH/gL/UL128-131, are known to play crucial roles in host cell entry (4–6). These two complexes share the same gH/gL heterodimer forming either with gO or with UL128 a disulfide bridge with gL-Cys144 (7). Cell-free virions which are infectious for multiple cell types rather than fibroblasts alone, thus resemble *in vivo* cell tropism, must harbour both gp complexes (8–10). HCMV strains show large differences in the relative levels of trimer and pentamer incorporated in their virions (11). It is suggested that the trimer–to-pentamer ratio influences the infection efficiency for the respective cell types (8, 12, 13) and that a number of HCMV genes have the capacity to impact the composition of the two gH/gL complexes (14). Large sequence comparison analysis has shown that, among all subunits of the two gp complexes, glycoprotein O (gO) exhibits by far the highest sequence polymorphisms with up to 23% amino acid diversity among gO sequences (15–17). All known gO sequences cluster into 5 major groups which can further be divided into 8 genotypes (18, 19). A closer inspection of gO gene sequences in circulating HCMV strains revealed that recombination among distinct strains may have occurred at several positions along the gO gene (16, 17, 20–22), arguing that recombination may be an important driving force of gO sequence evolution. Although it appears that all 8 gO genotypes can form stable trimers (11), it is poorly understood what role gO polymorphism plays in cell tropism. As recently shown gO genotypes may influence the efficiency of epithelial cell infection through specific sequence characteristics (23) or via affecting the relative levels of gH/gL complexes (11, 13). Moreover, it has recently been reported that the accessibility of certain gH or gH/gL epitopes for monoclonal antibodies differs among HCMV strains probably due to the distinct gO genotype sequences of the respective strains (24).

Over the last few years, a number of cellular interaction partners for both, the trimer and the pentamer have been identified (14). One of these cellular receptors, platelet-derived growth factor receptor alpha (PDGFRα), was identified to directly and specifically interact with gO parts of the trimer (25–27). This interaction enables entry of cell-free virions into fibroblasts, the only cell type which shows a high PDGFRα expression (28). Albeit, soluble forms of PDGFRα (sPDGFRα) can severely inhibit not only entry into fibroblasts, but also entry into endothelial and epithelial cells (25–27), and first observations indicate that the inhibitory capacity of sPDGFRα is effective against several HCMV strains even when they harbour a different gO genotype sequence (26).

Neuropilin-2 (NRP2), another recently identified host cell receptor for HCMV, specifically interacts with the UL128 subunit of the pentamer (29). This interaction is required for entry into endothelial and epithelial cells, most likely through endocytosis, but seems to be dispensable for entry into fibroblasts. Accordingly, a soluble form of NRP2 (sNRP2) inhibits endothelial and epithelial infection but not fibroblasts (29). Both PDGFRα and NRP2 likely function as the primary entry receptors for the trimer and pentamer, respectively, however, the modes of entry downstream of receptor binding may substantially differ. In particular, it appears that the trimer functions at steps which are required for entry into all cell types (14) which makes sPDGFRα or derivatives thereof a promising therapeutic tool against HCMV (30, 31). In the present study, we now aimed to assess how gO polymorphism influences the inhibitory capacity of sPDGFRα and sNRP2, respectively. To this end, we generated a set of TB40-BAC4-luc-derived HCMV gO genotype mutant strains, five of them harbour one of the major gO genotype sequences and 2 of them carry a recombinant gO genotypic form. We showed that subtle to moderate differences in the inhibitory capacities of the two entry inhibitors, sPDGFRα and sNRP2, are attributed to gO polymorphism.

## Results

### Cell-free infectivity of HCMV strains upon swapping of gO genotype sequences

In order to investigate the influence of gO polymorphisms on the cell entry inhibitors sPDGFRα and sNRP2, we generated a panel of gO genotype mutant viruses, in which the parental gO genotype sequence GT1c of TB40-BAC4-luc was fully or partially replaced by another gO GT sequence (see Figure 1, and Supplementary Figure 1). Correctness of the whole UL and US regions of fibroblast-derived reconstituted viruses were validated by whole genome sequencing. All experiments were done with reconstituted virus stocks without further passaging. For comparison analyses and comprehensiveness both the parental strain, referred to as gO GT1c, and the previously generated gO GT mutant, gO GT4 (23), were included in all experiments. First, to assess the ability of the gO GT mutants to infect human foreskin fibroblasts (HFFs), cell-free virus stocks of gO GT1c and mutants were adjusted to a similar number of encapsidated genome equivalents (mean of 8.2 log_10_ copies/ml). Infectivity was quantified by monitoring luciferase expression in cell lysates 2 days post infection. Relative light units (RLUs) are the read out for the extent of infection. The log_10_ ratio of RLUs to encapsidated genomes was calculated and the fold change relative to gO GT1c was determined. Mutants and parental strain were incubated on the same plate to avoid inter-plate variability of RLU quantitation. As shown in Figure 2A, all gO GT mutants infected fibroblasts similarly efficient as gO GT1c.

**Figure 1.**
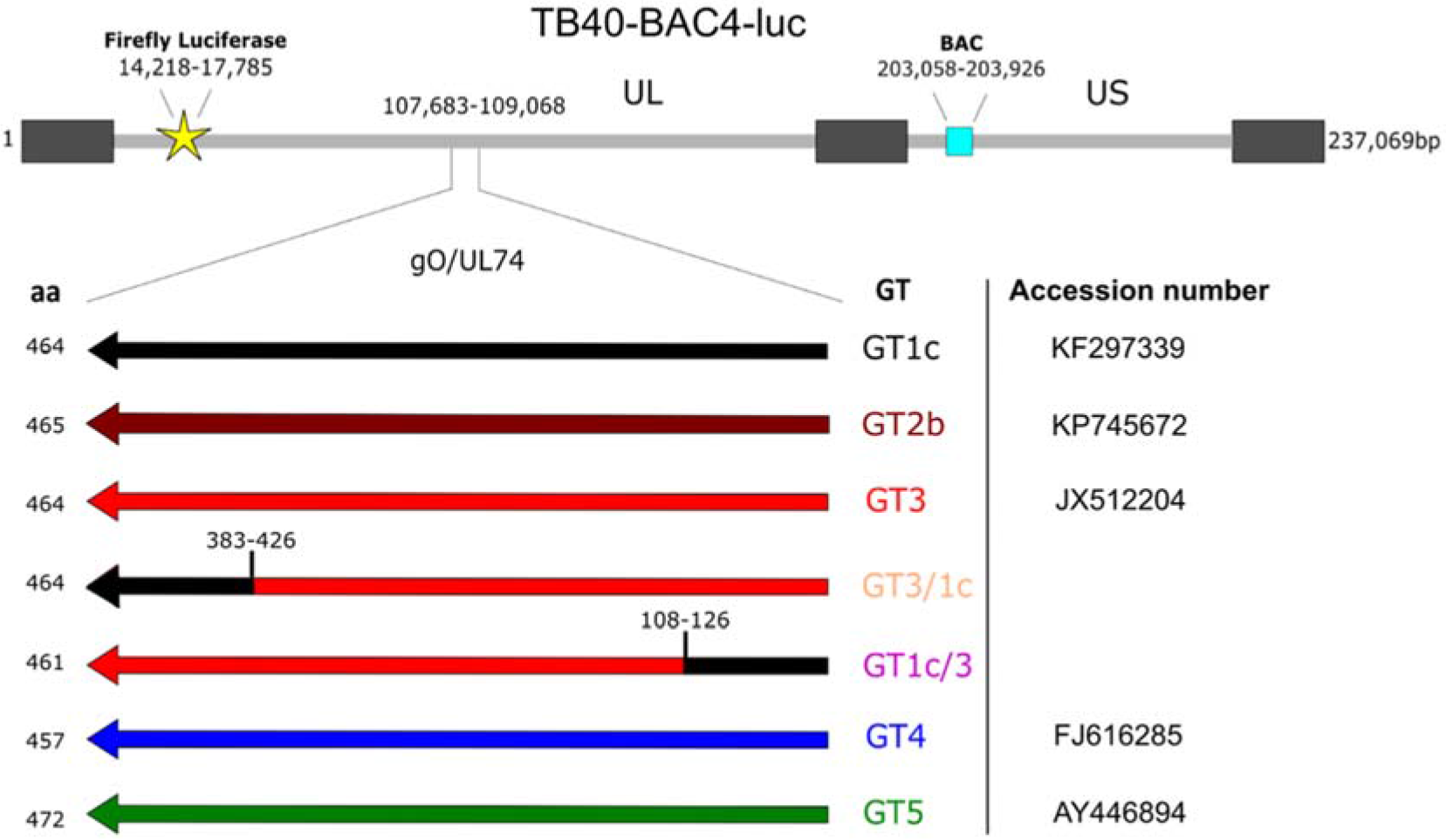
Schematic illustration of BAC-derived gO genotype mutants. The resident gO genotype (GT) 1c sequence of parental strain TB40-BAC4-luc was fully or partially replaced by the indicated gO GT sequences via “en passant” mutagenesis. Main genome characteristics are displayed. Arrows represent orientation and position of gO ORFs upon GT swapping. GTs and accession numbers of the HCMV strains from which the respective gO GT sequences are derived are shown on the right and the length of gO amino acid (aa) sequence on the left. Aa range of the recombination breakpoint of the chimeric mutants GT3/1c and GT1c/3 are depicted above the ORF. Cell-free bacterial artificial chromosome-derived mutant virus stocks were generated upon reconstitution in human foreskin fibroblasts.

**Figure 2.**
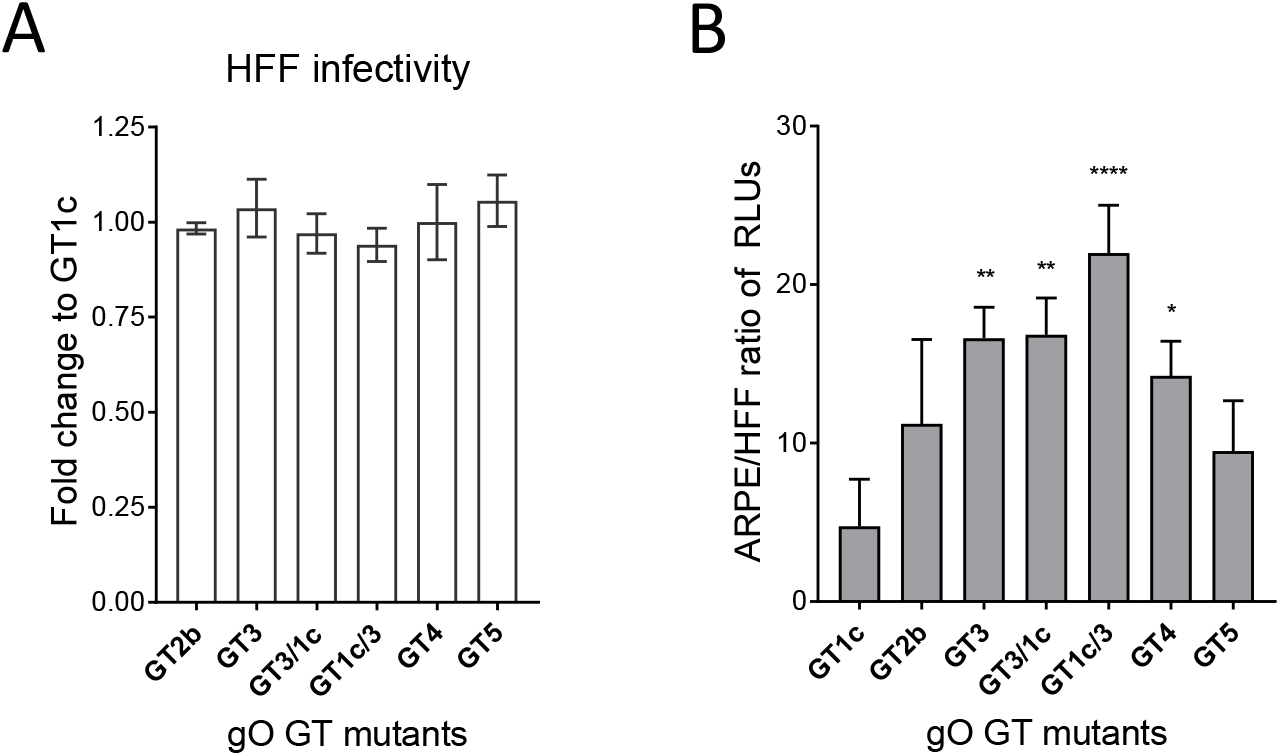
Cell-free infectivity of gO genotype mutants for fibroblasts and epithelial cells. A) Human foreskin fibroblasts (HFFs) were infected with parental strain gO GT1c and the panel of gO GT mutants using similar numbers of encapsidated genome equivalents (range: 8.0 – 9.2 log_10_ copies/ml). Two days post infection relative light units (RLUs) were assessed in cell lysates by luciferase assay as a read out of infection efficiency. Log_10_ RLU to genome ratio was calculated and the fold change relative to gO GT1c was determined. All experiments were performed in triplicates and data shown are means ± SEM of 2 - 4 independent experiments. B) HFFs and ARPE-19 cells were simultaneously infected with parental strain gO GT1c and the gO GT mutants using the same virus preparation for both cell types. Two days post infection RLUs were determined and the ratio of fibroblast to epithelial cell RLUs was calculated. All experiments were performed in triplicates and data shown are means ± SEM of 3 - 4 independent experiments. Statistical significance was evaluated by ANOVA with Tukey’s test for multiple comparison. ****p<0.0001; **p<0.01; *p<0.05 in comparison to GT1c.

Next, we determined the relative epithelial cell infectivity by simultaneously infecting both, epithelial cells and fibroblasts. Cell-free virus stocks were adjusted to achieve 300-1,500 RLUs in ARPE-19-infected cell lysates. Infection efficiencies were determined by luciferase assay 2 days post infection and the ratios of epithelial cell to fibroblast RLUs were calculated (see Figure 2B). Mutant gO GT3 and the two recombinant forms, GT1c/GT3 and GT3/GT1c, along with gO GT4 displayed a significantly higher epithelial cell infectivity compared to gO GT1c. In summary, the data show that gO GT swapping, either full-length or partial, does not impair the capacity to infect fibroblasts but seems to affect epithelial cell tropism.

### Content of gO and gH in the envelope of gO GT mutant viruses

Cell-free virions of parental strain TB40-BAC4-luc are characterized by high gO abundance and low UL128 expression. This is thought to result in a high trimer-to-pentamer ratio associated with a low efficiency for epithelial cell infection (12). Thus, we wanted to know whether the enhanced epithelial cell infectivity of gO GT3, gO GT1c/GT3 and gO GT3/GT1c results from changes in the trimer-to-pentamer ratio upon gO GT swapping. To this end, parental and mutant virions were purified from fibroblast supernatant and the amounts of gO and gH were determined by semi-quantitative western blot under reducing conditions. The total amount of virions was normalized to gB and/or major capsid protein (MCP). The gO content represents the amount of trimer and the gH level is assumed to indicate the overall amount of trimer and pentamer in virions. One representative immunoblot for each mutant is given in Figure 3, and the estimated virions’ gO and gH contents are shown in Table 1. In comparison to gO GT1c the amount of virions’ gO was severely reduced in gO GT3 (~ 80%), GT3/1c (~ 90%), GT1c/3 (~ 50%), a subtle reduction was found for gO GT5 (~ 30%), but no substantial changes for gO GT4 mutant virions. Notably, almost no gO was detectable in gO GT2b virions even when very high virion concentrations were used for immunoblotting (see Supplementary Figure 2). Although it cannot be excluded that gO GT2b virions harbour very low levels of gO it is more likely that the anti-gO antibody used in this study which is directed towards gO GT1c of TB40E (32) does not cross-react with gO GT2b while the gO genotypic forms GT3, GT4, and GT5 are well recognized (see Supplementary Figure 2). With regard to the gH content it appears that the mutant virions GT3 and GT1c/GT3 harbour 1.6 to 3.2 fold higher gH levels whereas GT2b, GT4, and GT5 contain moderately lower levels as compared to gO GT1c. Hence, these data suggest a shift towards lower trimer-to-pentamer ratio upon full-length and partial swapping of GT3 sequences but not upon replacement of GT1c by GT4 and GT5 sequences.

**Figure 3.**
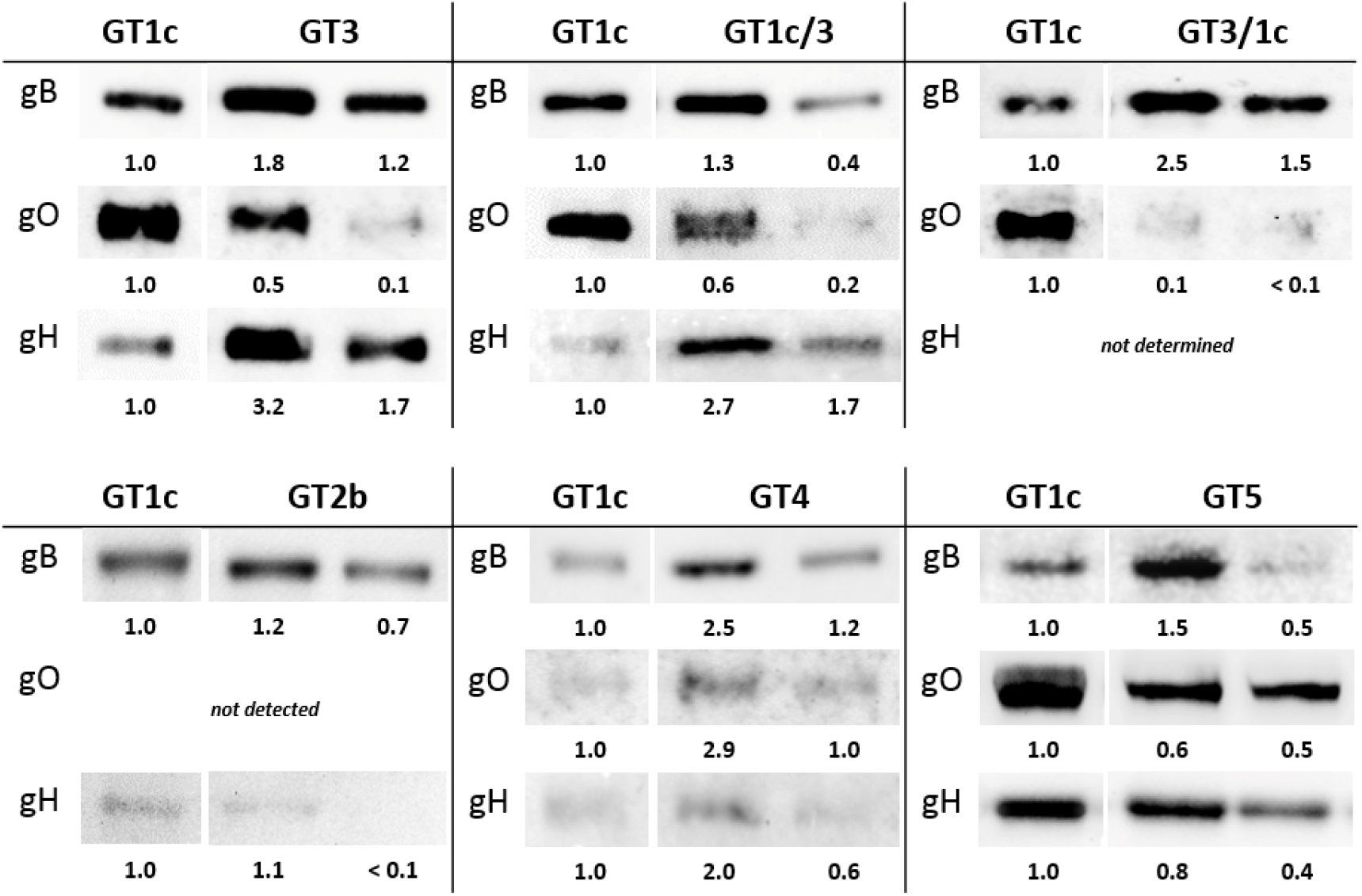
Comparison of gO and gH content in cell-free virions between parental strain gO GT1c and gO mutants. Virions harvested from human foreskin fibroblast supernatant were subjected to reducing gel electrophoresis and analyzed by Western Blot using antibodies directed against glycoproteins gB (anti-gB mAb 2F12), gO (anti-gO.02 mAb) and gH (AP86-SA4). The amount of virions loaded on the gels were compared to gB. Contents of gO and gH were compared between gO GT1c and the respective gO mutants. For each mutant an additional 2-fold dilution was loaded on the gel. Band densities were determined relative to the GT1c reference band for each blot individually, and are shown below the blots.

**Table 1.** Glycoprotein O and H content in fibroblast-derived cell-free virions.

| Name of gO<br>GT mutants | Envelope glycoproteins |  |
| --- | --- | --- |
|  | gO in % [mean (range)]* | gH in % [mean (range)]* |
| GT1c | 100 | 100 |
| GT2b | not detectable | 53 (14 - 92) |
| GT3 | 18 (8 - 28) | 160 (142 - 178) |
| GT3/1c | 6 (4 - 7) | not determined |
| GT1c/3 | 48 (46 - 50) | 317 (208 - 425) |
| GT4 | 100 (83 - 116) | 65 (50 - 80) |
| GT5 | 67 (53 - 80) | 70 (40 - 100) |

\*) normalized to envelope glycoprotein B

### Inhibition of cell-free fibroblast and epithelial cell infectivity by soluble PDGFRα-Fc

Since all gO GT mutants retained the ability to infect fibroblasts and epithelial cells, we were able to directly compare the inhibitory capacity of sPDGFRα-Fc between the five major gO genotypic and the two recombinant forms. The inhibition experiments were performed with a fixed amount of infectious viruses pre-incubated for 2 hours with a 2-fold dilution series of sPDGFRα-Fc ranging from 0.0025 to 0.625 μg/ml. After another 2 hour-incubation on fibroblasts or epithelial cells, respectively, cells were washed and subsequently incubated with fresh medium for further 2 days. RLUs were monitored by a luciferase assay and plotted against the concentration of sPDGFRα-Fc.

First, the appropriate amount of infectious input virus was determined using three different virus dilutions of parental strain gO GT1c. As shown in Figure 4A and 4B there was no substantial change in the overall shape of the dose-dependent inhibition over a wide range of input infectivity. Thus, for all further inhibition experiments the cell-free virus stocks were normalized to similar RLUs within the tested range (see Materials and Methods). In all mutants, cell-free infectivity was inhibited by sPDGFRα-Fc in a dose-dependent manner. One representative curve for gO GT1c and the gO GT mutants is shown in Figure 4C and 4D. The half-maximal inhibition (IC_50_) as calculated by non-linear regression ranged from 49 ng/ml to 73 ng/ml for fibroblasts and from 24 ng/ml to 56 ng/ml for epithelial cells (see Table 2). None of the mutants’ IC_50_ value significantly differed from IC_50_ of parental strain. Moreover, there was no difference between parental strain and mutants in the overall steep shape of the dose-response curves (slopes >1), neither in fibroblasts nor in epithelial cells, except for one of the recombinant mutants, gO GT1c/3, in epithelial cells. This gO mutant showed a shallower dose-response curve with a slope of 1.0 to 2.6 (see Table 2). The slope parameter mathematically analogous to the Hill coefficient is a measure of cooperativity (33) in the binding of multiple ligands (e.g. sPDGFRα-Fc) to linked binding sites (e.g. gO). Dose-response curves with a slope of about 1.0 are indicative for non-cooperativity which means the ligand binds at each site independently. In contrast, steep curves with slopes much larger than 1.0 are thought to result from a form of positive cooperative effects upon ligand binding (33). Hence, these findings suggest that the presumed positive cooperativity is weakened when sPDGFRα-Fc binds to gO GT1c/3.

**Figure 4.**
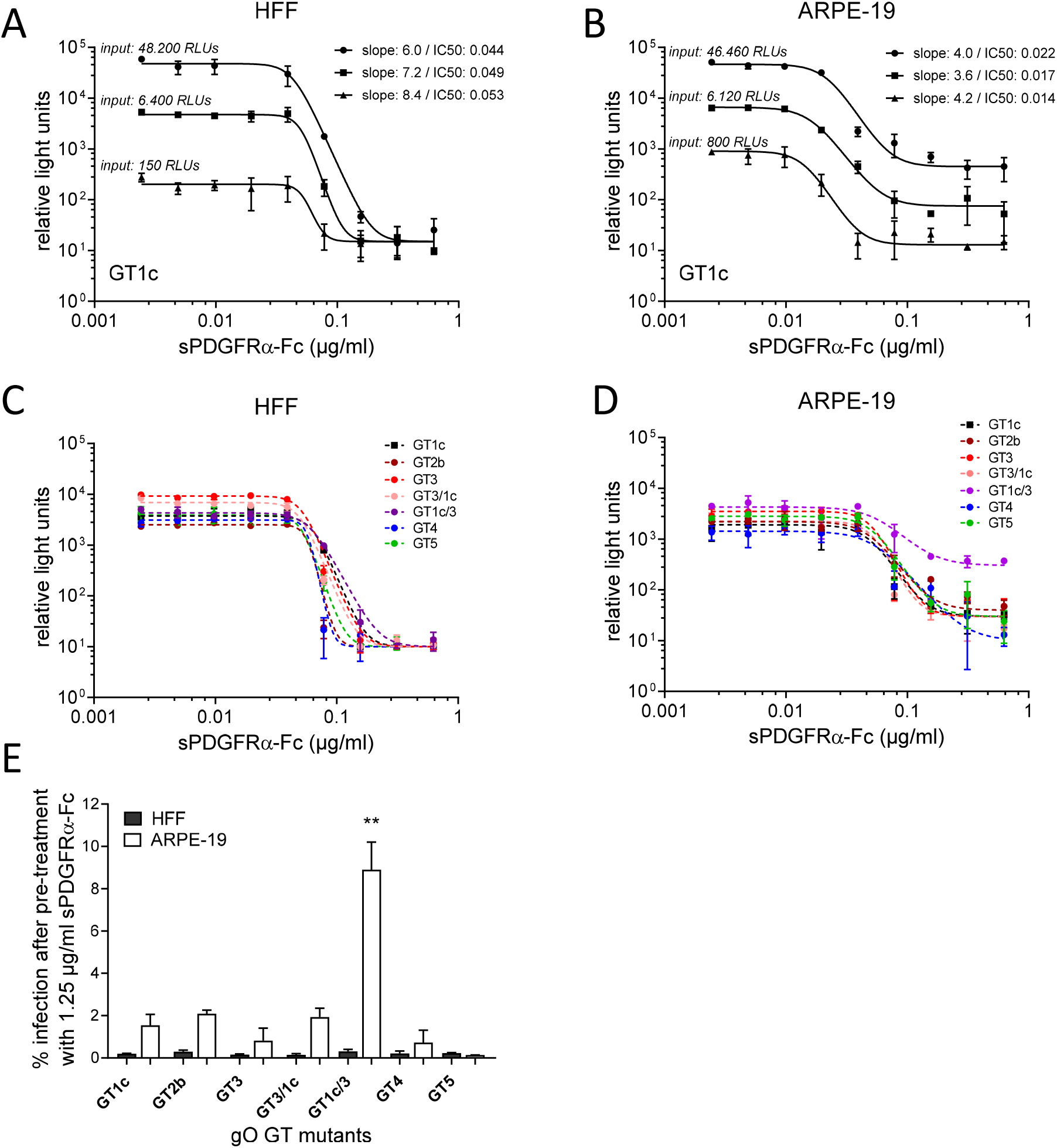
Inhibition of cell-free infectivity of gO genotype mutant viruses by soluble PDGFRα-Fc. The whole panel of gO genotype (GT) mutants along with parental strain gO GT1c were pre-incubated with soluble PDGFRalpha-Fc (sPDGFRα-Fc) before infection of human foreskin fibroblasts (HFFs) or adult retinal pigment epithelial cells 19 (ARPE-19 cells), respectively. Two days after infection relative light units (RLUs) were determined in cell lysates by a luciferase assay. In (A) and (B) three different virus stock concentrations of parental strain gO GT1c indicated as input RLUs are used. In (C to E) mutant virus stocks were diluted to achieve RLUs ranging from 1.000 to 19.000 without treatment. In (A to D) parental and mutant virus stocks were treated with serial 2-fold dilutions of sPDGFRα-Fc (range: 0.625 to 0.0244 μg/ml). Monitored RLUs were plotted against sPDGFRalpha-Fc concentrations. Four-parameter dose-response curves were generated and the protein concentration causing inhibition of 50% of infection (IC_50_) and the steepness of the curves were calculated (see in A and B and in Table 1). In (C) and (D) one representative curve from each mutant out of 2 – 4 independent experiments is shown. Data represent mean values ± SDs of triplicate determinations. In (E) the % of infection of HFFs or ARPE-19 cells, respectively, after pre-treatment with 1.25μg/ml sPDGFRα-Fc is shown. Experiments were performed in triplicates and data are means ± SEM of 3 - 5 independent experiments. Statistical significance was evaluated by ANOVA with Tukey’s test for multiple comparison. **p<0.01 in comparison to GT1c.

**Table 2.** Dose-inhibition curve characteristics of gO genotype mutant strains.

| Name of gO<br>GT mutants | sPDGFRα-Fc dose-response<br>curves in HFFs | | sPDGFRα-Fc dose-response<br>curves in ARPE-19 cells | | sNRP2-Fc dose-response<br>curves in ARPE-19 cells | |
| --- | --- | --- | --- | --- | --- | --- |
|  | IC<sub>50</sub> in ng/ml<br>[mean (range)] | slope (range) | IC<sub>50</sub> in ng/ml<br>[mean (range)] | slope (range) | IC<sub>50</sub> in ng/ml<br>[mean (range)] | slope (range) |
| GT1c | 58 (52 - 63) | 6.0 - 8.0 | 39 (16 - 53) | 2.6 - 6.0 | 31 (18 - 44) | 0.9 - 1.1 |
| GT2b | 73 (54 - 92) | 6.0 - 8.0 | 36 (10 - 51) | 2.6 - 3.0 | 65 (22 - 110) | 1.2 - 1.5 |
| GT3 | 65 (53 - 82) | 4.8 - 10.0 | 47 (43 - 50) | 4.7 - 6.0 | 34 (27 - 40) | 1.1 - 1.5 |
| GT3/1c | 49 (46 - 54) | 6.0 - 9.5 | 50 (49 - 50) | 6.8 - 7.4 | 80 (50 - 110) | 0.9 - 2.1 |
| GT1c/3 | 69 (56 - 81) | 2.5 - 6.2 | 30 (13 - 62) | 1.0 - 2.6 | 65 (34 - 100) | 0.9 - 1.3 |
| GT4 | 52 (47 - 55) | 5.5 - 6.0 | 55 (52 - 56) | 5.3 - 6.0 | 90 (60 - 110) | 0.7 - 1.3 |
| GT5 | 68 (45 - 100) | 5.3 - 8.0 | 56 (54 - 58) | 5.3 - 6.0 | 67 (24 - 110) | 1.2 - 1.7 |

The data shown are the results of two to five independent experiments. IC<sub>50</sub> and slope values were calculated from [inhibitor] vs. response four-parameter dose-response curves.

Next, we determined the maximal extent of inhibition at 1.25 μg/ml sPDGFRα-Fc calculated as 1 – (RLU after pretreatment / RLU of untreated virus stocks). The inhibition of fibroblast infectivity was almost complete (> 99%) and did not differ between gO GT1c and mutants (see Figure 4E). In epithelial cells, in contrast, one of the recombinant mutants, gO GT1c/3, retained a significantly higher infectivity (mean: 9%) at this inhibitor concentration compared to gO GT1c. The other mutants did not differ from the parental strain. The reduced epithelial cell inhibition of gO GT1c/3 by sPDGFRα-Fc is well in accordance with the shallower shape of the dose-response curve (Figure 4D). Notably, the inhibition efficiency was slightly less effective in epithelial cells (~ 98 – 99%) than in fibroblasts for gO GT1c and the mutants, GT2b, GT3, and GT3/1c (see Figure 4E).

In summary, these findings show that not only the five major genotypic forms of gO are recognized by sPDGFRα-Fc but also recombinant forms of gO. Albeit, one recombinant version of gO seems to be less effectively inhibited by sPDGFRα-Fc on epithelial cells.

### Inhibition of cell-free fibroblast and epithelial cell infectivity by soluble NRP2-Fc

It has recently been reported that soluble forms of NRP2 which specifically bind to UL128, inhibit epithelial cell infection while fibroblast infection remains largely unaffected (29). We wanted to know whether alterations in the virions’ gO and gH content upon gO GT swapping may indirectly affect the inhibitory capacity of sNRP2-Fc. To address this question, first we performed inhibition experiments on epithelial cells using a 2-fold dilution series of sNRP2-Fc (range: 0.0025 to 0.626 μg/ml) and a fixed amount of RLU-normalized gO GT virions as described for sPDGFRα-Fc. Two independent experiments were performed for each mutant along with the parental strain and one representative curve is shown in Figure 5A. The dose-dependent inhibition was similar between parental strain and gO GT mutants, the dose-response curves displayed slopes of about 1.0 (range: 0.7 to 2.1) and the IC_50_ values ranged from 31 to 90 ng/ml (see Table 2). These findings indicate that neither the gO genotypic form nor changes in the virion’s gO content influences the capacity of sNRP2-Fc for epithelial cell inhibition. Finally, we wanted to assess the maximum inhibitory capacity of 1.25 μg/ml sNRP2-Fc on epithelial cells and whether such high inhibitor concentrations also have an effect on fibroblast infectivity. As shown in Figure 5B, epithelial cell infectivity was 96% to 98% reduced in all mutant viruses and this did not significantly differ between parental strain and mutants. Interestingly, although the fibroblast infectivity was almost unaffected in parental strain and in two of the mutants, GT2b and GT5, a moderate reduction in fibroblast infectivity of 20% to 40% was observed for the other mutants and this reached statistical significance for the recombinant mutant gO GT1c/3. From these data it appears that gO differences upon GT swapping may render mutant virions partially accessible to sNRP2-Fc inhibition on fibroblasts.

**Figure 5.**
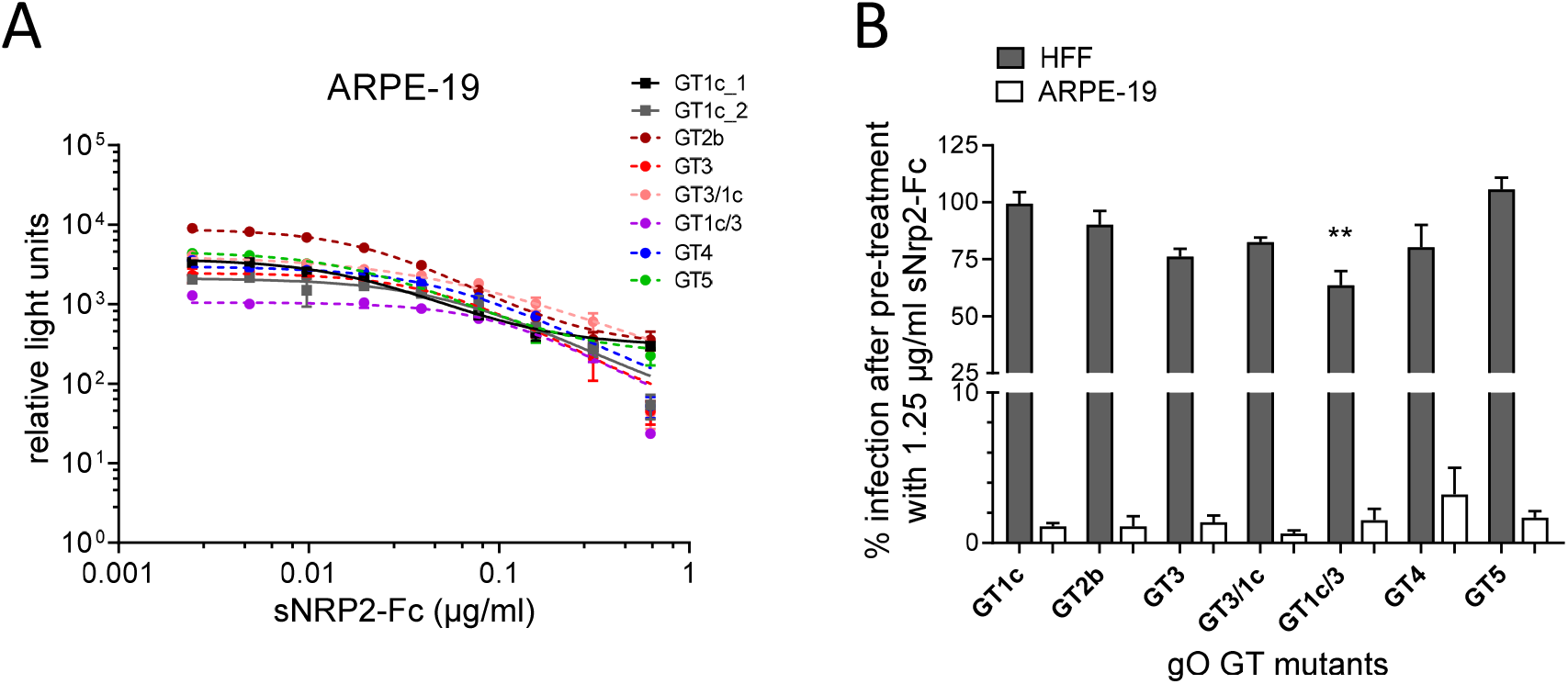
Inhibition of cell-free infectivity of gO genotype mutant viruses by soluble NRP2-Fc. A) Parental gO GT1c and gO GT mutant virus stocks were pre-incubated with serial 2-fold dilutions of soluble NRP2-Fc (sNRP2-Fc) (range: 0.625 to 0.0244 μg/ml) before infection of adult retinal pigment epithelial cells 19 (ARPE-19 cells). Two days after infection relative light units (RLUs) were monitored ranging from 1.000 to 10.000 RLUs in untreated controls. RLUs were plotted against sNRP2-Fc concentrations and four-parameter dose-response curves were generated to calculate the protein concentration causing inhibition of 50% of infection (IC50) and to determine the steepness of the curves. Two curves from gO GT1c and one representative curve from each mutant out of 2 independent experiments is shown. Virus stock concentrations used were similar as for (A). Data represent mean values ± SDs of triplicate determinations. B) The % of infection of HFFs or ARPE-19 cells, respectively, after pre-treatment with 1.25μg/ml sNRP2-Fc is shown. Experiments were performed in triplicates and data are means ± SEM of 3 - 5 independent experiments. Statistical significance was evaluated by ANOVA with Tukey’s test for multiple comparison. **p<0.01 in comparison to GT1c.

## Discussion

The two envelope glycoprotein complexes, gH/gL/gO-trimer and gH/gL/UL128L-pentamer, which share the same gH/gL heterodimer, play major roles in HCMV cell entry. In the present study, we focussed on gO, the critical subunit of the trimer. A special hallmark of gO is its high polymorphism with an overall amino acid diversity of ~ 20% (18, 19). To learn more about potential functional differences attributed to gO polymorphism, we fully or partially swapped gO gene sequences in the otherwise identical TB40-BAC4-luc background, tested the set of gO mutants for their capability to infect fibroblasts and epithelial cells, for their relative composition of gO and gH in cell-free virions, and evaluated the inhibitory capacity of sPDGFRα-Fc in comparison to sNRP2 inhibition.

First, we demonstrate that gO GT swapping, either partial or full-length, does not substantially affect fibroblast infectivity but may lead to an increase in relative epithelial cell infectivity. In particular, the mutants gO GT3, GT3/1c, and GT1c/3, which displayed the strongest enhancement in epithelial cell infection, contained substantially lower gO but higher gH levels in their cell-free virions as compared to parental strain. Previous studies have revealed that gO and UL128 compete for binding to the same gL cysteine residue in gH/gL (7) which in turn regulates the trimer to pentamer ratio (7, 8) and this renders virions more infectious for fibroblasts (high trimer to pentamer ratio) or epithelial cells (low trimer to pentamer ratio) (11, 12). Since parental strain TB40-BAC4-luc is characterized by vastly more gO than UL128 accompanied by a low epithelial cell infectivity (12) it is likely that the opposite changes in gO and gH levels upon partial or full-length GT3 swapping cause a shift towards lower trimer to pentamer ratio which may well explain the increase in epithelial cell infectivity. The impact of the relative composition of gO and gH in terms of epithelial cell infectivity is further underlined by the observation that replacement of gO GT1c by GT5 which causes a subtle reduction in both, gO and gH, has no effect on epithelial cell infectivity. Similarly as recently reported, gO GT1b to GT5 swapping and vice versa has also no effect on gO expression levels (13). Together, these findings suggest that the relative abundance of gO and gH incorporated into cell-free virions is influenced by the gO genotypic form. Recently, UL148 has been identified to regulate the trimer to pentamer ratio by stabilizing gO within the endoplasmatic reticulum (34, 35). It is tempting to speculate that the regulatory capability of UL148 is influenced by the gO sequence. Additionally, it cannot be excluded that GT sequence-specific characteristics directly modify the capacity of cell-free virions for entry into epithelial cells, since gO GT4 displayed an enhanced epithelial cell tropism without substantial alterations in gO and gH abundance, as previously shown (23).

Remarkably, despite significant differences in epithelial cell tropism, the fibroblast infectivity was similar among the mutants and parental strain. These findings indicate that neither changes in gO and gH abundance nor gO GT sequence-specific characteristics affect the capacity for fibroblast infection. Moreover, these data lead to the conclusion that all of these gO genotypic forms can bind to the cellular fibroblast receptor PDGFRα with similar efficiency. Strikingly, a minimum level of gO on cell-free virions seems to be still sufficient for normal fibroblast infection under the tested *in vitro* conditions as in particular gO GT3 and gO GT1c/3 mutants display very low gO levels. Taken together, these observations provide clear evidence that gO polymorphism has a substantial impact on epithelial cell but not on fibroblast infectivity. Further investigations will clarify how differences in the gO and gH abundance and/or GT-specific sequence characteristics affect epithelial cell entry of cell-free virions.

Recombination among different HCMV strains appears to be a major driving force in HCMV evolution as shown by numerous studies (36). Recently, the recombination density throughout the genome was deeply investigated by whole genome sequence comparisons exploring past and recent recombination events as well (16, 20, 21). A particularly interesting finding was the identification of pervasive genome-wide recombination generating diversity both within and between genes (16, 21). So far, little is known about potential functional consequences for individual genes upon intragenic recombination. In the present study, we have now included two chimeric gO GT mutants, each of them carrying a recombinant gO genotypic form composed of GT1c and GT3 sequences. One of them, gO GT3/1c, harbours the recombination breakpoint within the conserved C-terminal part of gO and this mutant differs from full-length gO GT3 in only 4 amino acid residues. The recombination breakpoint of the other one, gO GT1c/3, is located in a small identical sequence stretch between GT1c and GT3 in the otherwise highly polymorphic N-terminal part of gO. Recombination resulted in a severely altered gO sequence with an amino acid diversity of 9% from GT1c and of 10% from GT3. Strikingly, both recombinant gO mutants not only fully retained the ability to infect fibroblasts they even displayed an enhancement in epithelial cell infectivity comparable to full-length GT3 mutant. As discussed above, the change in gO and gH abundance may cause the observed epithelial cell phenotype. Accordingly, these findings indicate that recombination within the gO gene could be considered as an important function for HCMV to generate (i) gene diversity with or without modified functions and (ii) novel combinations of neighbouring loci even when they are highly diverse. This is well in concordance with previously reported sequencing data showing that recombination within gO may sporadically occur also *in vivo* despite a strong linkage between gO and the adjacent, partly overlapping gN gene (16, 17, 21, 22, 37).

Recent studies have shown that PDGFRα specifically interacts with the gO subunit of the trimer which is required for entry into fibroblasts (25–28). As soluble forms of PDGFRα or derivatives thereof can inhibit cell-free infection of several cell types (26) it appears that binding of sPDGFRα to gO interferes with trimer-mediated function(s) widely required for cell entry. We now demonstrate that representatives of the five major gO genotypic forms, GT1c, GT2b, GT3, GT4, and GT5, are similarly recognized by sPDGFRα and upon pretreatment with sPDGFRα-Fc both, fibroblast and epithelial cell infectivity was strongly inhibited. These data are well in line with previous reports showing the inhibitory capacity of sPDGFRα for several distinct HCMV strains (26). Notably, even at a concentration of 1.25μg/ml sPDGFRα we observed a residual infectivity of about 1 - 2% in epithelial cells, while in fibroblasts the inhibition was almost complete (≥ 99%) similar as shown previously (27). Thus, it is tempting to speculate that a trimer-independent entry mechanism accounts for the residual infectivity. Alternatively, it is also possible that not all virions are neutralized at this concentration allowing for a residual infection.

Remarkably, one of the two recombinant mutants, gO GT1c/GT3, displayed a significantly lower sensitivity for sPDGFRα inhibition on epithelial cells than the other mutants while the fibroblast inhibition was similarly effective. As mentioned above this mutant comprises its recombination site in the highly polymorphic N-terminal region of the protein, which only recently was suggested to contain the PDGFRα receptor binding domain (31). By mutational analysis the authors identified a small stretch from amino acid 117 to 121 causing the strongest impairment of sPDGFRα binding to virus particles and consequently also a reduced virus penetration into fibroblasts. Although this peptide site overlaps with the recombination site of GT1c/GT3, the specific sequence remained unchanged upon recombination suggesting that sPDGFRα binding to this recombinant form of gO is not impaired. This presumption fits well to the finding that gO GT1c/GT3 mutant does not display a phenotype in fibroblast infectivity while mutants with a mutation in this particular peptide sequence showed reduced penetration into fibroblasts (31). Hence, we assume that the impaired sPDGFRα inhibition for gO GT1c/GT3 mutant on epithelial cells is not caused by a lower binding of sPDGFRα to gO but rather by an impaired interference with a downstream entry step mediated by the trimer.

The assumption that binding of sPDGFRα to gO-trimer affects more entry properties than the unique block of the receptor binding site is further strengthened by our findings that inhibition with 2-fold serial dilutions of sPDGFRα led to steep dose-response curves in both fibroblasts and epithelial cells. Such steep inhibition curves with a slope of much greater than 1 are thought to result from a form of positive cooperative effects upon ligand binding (33). Remarkably, the steepness of the sPDGFRα-Fc dose-inhibition curves were not affected by the gO content in virions, nor by the amount of input virions. The underlying mechanisms are not yet clearly understood but following scenarios may explain why sPDGFRα-bound virions become rapidly inactive for cell entry: binding of sPDGFRα to virions leads to (i) steric hindrance and/or conformational changes of the gO-trimer which affects multiple sPDGFRα binding sites on the virion, (ii) cluster formation of trimers and/or other envelope complexes which causes that multiple gO binding sites on the virion are rapidly blocked, (iii) changes of gB prefusion into gB postfusion conformation (under the assumption that the trimer stabilizes the gB prefusion conformation) which renders virions inactive for entry, and/or (iv) cluster formation of multiple virions. Although these proposed scenarios await further clarification, from our data it becomes clear that a presumed cooperative effect triggered by sPDGFRα does not differ among the five major gO genotypes.

When we compared the dose-response curves of sPDGFRα with those of sNRP2, a recently identified entry inhibitor for epithelial cells (29), it becomes obvious that the mechanisms of action substantially differ between these two entry inhibitors. The dose-response curves of sNRP2 displayed a slope of about 1 – 2 meaning that binding of sNRP2 to its interaction partner UL128 of the pentamer causes no further effects beside the block of the binding site. There was also seen no difference among the gO GT mutants and parental strain indicating that neither gO abundance nor gO GT-specific characteristics influence the binding efficiency of sNRP2. Notably, all gO mutants along with parental strain displayed a residual epithelial cell infectivity of about 2-3% at a concentration of 1.25 μg/ml sNRP2. Whether an NRP2-independent entry pathway circumvents a complete inhibition or whether not all virions are neutralized by this concentration of sNRP2 yet awaits further investigation. As recently reported, fibroblast infection is largely unaffected by sNRP2 (29). In overall, this finding is well in concordance with our data. However, we observed a subtle inhibition of fibroblast infectivity by high concentrations of sNRP2 in those mutant virions which displayed very low amounts of gO. Thus, it is presumable that binding of high amounts of sNRP2 to virions lead to steric hindrance of the trimer which becomes visible only for virions with low gO levels.

In conclusion, in this study we show that the trimer to pentamer ratio which is substantially affected by gO polymorphism has no influence on the inhibitory capacity of sPDGFRα but may render virions slightly susceptible to sNRP2 inhibition on fibroblasts. When sPDGFRα or derivates thereof are considered for a therapeutic option to HCMV infection it should be taken into account that gO intragenic recombination may lead to partial evasion from sPDGFRα inhibition.

## Material and Methods

### Cells

Human foreskin fibroblasts (HFFs) were cultured in minimum essential medium Eagle (MEM; Sigma-Aldrich, St. Louis, Missouri) supplemented with 10% heat-inactivated fetal bovine serum (FBS; Capricorn Scientific, Ebsdorfergrund, Germany) and 0.5% neomycin (Sigma-Aldrich). Human adult retinal pigmented epithelial cells (ARPE-19; ATCC, Manassas, Virginia) were cultured in Dulbecco’s modified Eagle medium/nutrient mixture F12 (PAN-Biotech, Aidenbach, Germany) supplemented with 10% FBS and 1% penicillin-streptomycin (Thermo Fisher) or in MEM supplemented with 10% FBS and 0.5% neomycin.

### Generation of gO mutant BAC clones by en passant mutagenesis

All HCMV gO mutant strains were derived from the bacterial artificial chromosome (BAC) clone TB40-BAC4-luc (38). By “en passant” mutagenesis in *E.coli* GS1783 (39), the gO GT1c sequence of TB40-BAC4-luc was fully exchanged by GT2b, GT3, and GT5 respectively, and partially by gO GT3, either at the 5’ or 3’ end of gO GT1c ORF. For generation of full-length gO BAC mutants, a gO deletion mutant was used in which the whole gO ORF sequence was deleted. This ensured recombination between transfer plasmid and BAC-DNA solely upstream and downstream of the gO ORF sequence. For generation of recombinant BAC mutants, GT3/1c and GT1c/3, original TB40-BAC4-luc BAC DNA was used and both chimeric versions resulted from recombination within the gO GT1c ORF. First, a set of recombination cassettes were generated and the primer pairs used are listed in Supplementary table 1. For this, inserts containing a kanamycin resistance gene, flanked on one side by an 18-bp I-Sce I restriction sequence and a gO GT-specific 50-bp sequence, and on both sides by a Sac I or Nde I restriction site, respectively, were generated by PCR using pEP-Kan-S (kindly provided by Nikolaus Osterrieder). Second, each individual insert was cloned into the corresponding restriction site of gO GT sequence carried by pEX-A258 ordered from Eurofins Genomics (Luxembourg). The resulting transfer plasmids were used as template to generate the PCR-derived recombination cassettes containing extensions of ~50 bp sequences on each end for homologous recombination. The recombination cassettes were electroporated into recombination-competent *E.coli* GS1783 carrying the full-length or gO-deleted TB40-BAC4-luc DNA. After electroporation, recombination-positive *E.colis* were subjected to kanamycin selection, and the introduced non-HCMV sequences were removed within *E.coli* by cleavage at the I-Sce I site and a second red recombination. Positive kanamycin-sensitive, chloramphenicol-resistant bacteria colonies were selected. Finally, recombinant BAC DNAs were isolated from positive clones and the correctness of the BAC DNA sequence was verified by whole genome sequencing (see below). Further, overnight *E.coli* cultures of positive clones were stored at −80 °C until further use.

### BAC-derived gO mutant HCMV strains

Infectious viruses were generated by reconstitution as described previously (23). Briefly, mutant BAC DNAs were purified from *E.coli* using the Nucleobond BAC100 kit (Macherey-Nagel, Düren, Germany). The day before transfection, HFFs were seeded in 6-well plates (3 x 10^5^ cells/well) and then 2 μg of BAC DNA, 1 μg of pCMV71 DNA (plasmid was kindly provided by Mark Stinski, University of Iowa) and 9 μl of ViaFect reagent (Promega, Madison, Wisconsin) were mixed together with 100 μl of MEM without antibiotics, incubated for 15 min at room temperature and then added to the cells. 24 h after transfection, cells were washed with PBS and fresh MEM with antibiotics was added. One week after transfection, cells were trypsinized and transferred into 25 cm^2^ cell culture flasks. When CPE was 90-100%, supernatants were cleared by centrifugation at 4°C for 20 min at 4,000 x *g* and stored as cell-free viral stocks in aliquots at −80 °C. For infection and inhibition analyses, all aliquots were used only once to avoid multiple freeze-thaw cycles. Furthermore, one aliquot per reconstitution was subjected (i) to next generation sequencing to confirm the correctness of the complete UL and US genomic regions, (ii) to DNase treatment to assess the amount of encapsidated genomes, and (iii) to RLU measurements in order to normalize virus stocks in subsequent experiments. Two independent reconstitutions were performed for each mutant.

### Whole genome sequencing

DNA from BAC purification (as described above) and extracted DNA from DNase-treated or untreated viral stocks from HFF cell culture supernatants upon reconstitution were quantified using the Qubit 2.0 fluorometer (Thermo Fisher) according to the manufacturer’s instructions. One to two ng of DNA per sample were taken for library preparation using the Nextera XT DNA Library Preparation Kit and uniquely indexed samples using the Nextera XT Index Kit were pooled and sequenced together (both Illumina, San Diego, California). Pooled libraries were sequenced with paired-end reads (2×150-250) on a MiSeq system using v2 or v3 sequencing reaction chemistry (Illumina). Data were analyzed by CLC genomics workbench v12 software (Qiagen). Low-quality reads were trimmed and in average 52 - 80% of reads mapped to the reference genome.

### Determination of encapsidated HCMV genomes in virus stocks

In order to remove non-encapsidated viral DNA and free cellular DNA, fibroblast-derived virus stocks were treated with TurboDNase (Thermo Fisher). For this, 100 μl of master mix (73 μl H_2_O, 20 μl 10x DNase buffer, 5 μl 10x PBS, 2 μl TurboDNase (2 u/μl)) were added to 100 μl of sample and incubated for 1 h at 37°C in a thermoshaker at 1,400 rpm. Immediately thereafter, the total reaction volume was added to 2ml lysis buffer and DNA was extracted using the bead-based NucliSens EasyMag extractor (BioMérieux, Marcy-l’Étoile, France) according to the manufacturer’s protocol. DNA was eluted in 50 μl of nuclease-free H_2_O.

### HCMV-specific quantitative PCR

HCMV-DNA was quantitated using an in-house real-time qPCR amplifying a conserved region within US17 (forward primer GCGTGCTTTTTAGCCTCTGCA (10 pM), the reverse primer AAAAGTTTGTGCCCCAACGGTA (10 pM), TaqMan probe FAM-TGATCGGCGTTATCGCGTTCTTGATC-TAMRA (2 pM)) as previously described (23).

### Normalization of parental and mutant virus stocks

The firefly luciferase gene of HCMV strain TB40-BAC4-luc allows to monitor relative light units (RLUs) in infected cell lysates as a read out for infection efficiency (40). For normalization of parental strain and mutant virus stocks to similar RLUs, HFFs and ARPE-19 cells were seeded in white, clear, flat-bottom 96-well plates (Corning, Corning, New York) at a density of 1 x 10^4^ cells/well. The following day, viral stocks were serially 2-fold diluted in cell culture medium and 100 μl of viral dilution per well were used to infect the cells in triplicates for 2 h at 37 °C. Cells were washed three times with PBS, supplied with 100 μl of medium and incubated further at 37 °C for 2 days. RLUs were determined by luciferase assay of cell lysates according to the manufacturer’s protocol (SteadyGlo Luciferase Assay System, Promega) and measured in a Victor Light 1420 plate reader (PerkinElmer, Waltham, Massachusetts). Mean RLUs of triplicates were calculated for normalization. For inhibition assays viral stock dilutions generating 1,000 to 20,000 RLUs for both, HFFs and ARPE-19 cells were used. For determination of relative epithelial cell infectivity viral stock dilutions generating 300-1,500 RLUs in ARPE-19 cells were used.

### Fibroblast infection efficiency

HFFs were seeded in white, clear, flat-bottom 96-well plates (Corning, Corning, New York) at a density of 1 x 10^4^ cells/well. The following day, viral stocks were diluted to similar number of encapsidated genomes (range: 8.2 to 9.2 log_10_ genome copies/ml) in cell culture medium as previously determined and 100 μl of viral dilution per well were used to infect the cells in triplicates for 2 h at 37 °C. Cells were washed three times with PBS, supplied with 100 μl of medium and incubated further at 37 °C for 2 days before monitoring mean RLUs of technical triplicates. In parallel, 5 μl of the viral dilution was used to determine the actual number of encapsidated genomes used for infection. The ratio of log_10_ RLUs to log_10_ encapsidated genomes of parental strain was set at 1.0 and the fold change of the mutants as compared to parental strain was calculated. Three to four independent experiments per mutant were performed.

### Relative epithelial cell infectivity

The same viral stock dilution was used for infection of both fibroblasts and epithelial cells each seeded in white, clear, flat-bottom 96-well plates at a density of 1 x 10^4^ cells/well one day before infection. Two days after infection, RLUs were determined by luciferase assay as mentioned above and the epithelial to fibroblast RLU ratio was calculated. All experiments were performed in technical triplicates and three to four independent experiments were performed.

### Production of purified virions for immunoblotting

Supernatants from infected HFFs were harvested when cells displayed > 90% CPE and then clarified by centrifugation at 4,000 x *g* for 30 min at 4 °C. After filtration through a 0.45 μm filter (Whatman, GE Healthcare Life Sciences, Thermo Fisher) viruses were concentrated by centrifugation at 4 °C using vivaspin 20 concentrators with a molecular weight cutoff of 100K (Sartorius, Göttingen, Germany). Thereafter, virions were purified by ultracentrifugation through a 20% sucrose TAN (0.05 M triethanolamine, 0.1 M NaCl, pH 8.0) cushion for 80 min at 70,000 x *g* at 4 °C and the pellets were gently resuspended in TAN buffer on ice and stored in aliquots at −80 °C until further use.

### Western blot analysis

For sample preparation, virus stocks were mixed undiluted or diluted in TAN buffer with an equal volume of reducing 2x sample buffer (125 mM Tris/Cl pH 6.8, 6% SDS, 10% glycerol, 10% 2-mercaptoethanol, 0.01% bromophenolblue) and incubated on ice for 10 min before boiling at 95 °C for 10 min. Samples were separated on 10% SDS PAGE gels together with a high-range rainbow marker (Amersham ECL High-Range Rainbow Molecular Weight Marker, GE Healthcare, UK). Separated proteins were transferred to polyvinylidene difluoride (PVDF) membranes (Immun-Blot, Bio-Rad, California, USA) in blotting buffer (40 mM Tris, 39 mM glycine, 1.3 mM SDS, 20% methanol), which were then incubated overnight in blocking buffer (PBS, 1% BSA, 0.1% Tween-20) at 4 °C. All antibodies (Abs) were diluted in blocking buffer. Primary mouse anti-gH (AP86-SA4) and anti-MCP monoclonal antibodies (mAbs), anti-gO.02 mAb, and gB antibody (2F12; Abcam, Cambridge, UK) were incubated for 2 h at RT. Sheep, anti-mouse IgG-HRP (Amersham, GE Healthcare, UK) was used as secondary antibody and incubated for 1 h at RT. SuperSignal West Femto Maximum Sensitivity substrate (Thermo Fisher) was applied for gO detection according to the manufacturer’s instructions and Pierce ECL Western Blotting Substrate (Thermo Fisher) for gH, MCP, and gB detection. Chemiluminescent signals were visualized and analyzed using the ChemiDoc Imager and the Image Lab 6.0 software (both Bio-Rad).

### Inhibition assays

Cells were seeded in white, clear, flat-bottom 96-well plates at a density of 1 x 10^4^ cells/well the day before infection. A fixed amount of virus as determined by RLU normalization was pre-incubated with serial dilutions of soluble forms of PDGFRα-Fc or NRP2-Fc, respectively, for 2 hours before infection. Two hours after infection cells were washed twice with 1x PBS, supplied with 100 μl medium per well and further incubated for 2 days before subjected to luciferase assay. Total RLUs and percentage relative to RLUs of mock-preincubated controls were calculated. Two to five independent experiments per mutant viruses were performed and all experiments were carried out in technical triplicates. Another independently reconstituted BAC-derived virus per mutant was used to confirm the results.

### Statistical analyses

To compare relative epithelial cell infectivity (Figure 2) and percentage of infection after pre-treatment with soluble entry inhibitors, sPDGFRα and sNRP2, respectively, (Figures 4 and 5) between gO GT1c and gO GT mutants one-way ANOVA and Tukey’ tests for multiple comparison were used. Mean RLU values from three to five independently repeated experiments were used for statistical analyses. *P* values < 0.05 were considered significant. GraphPad Prism version 7.01 was used for statistical analyses.

## Acknowledgements

We are very grateful to Michaela Binder, Sylvia Malik, Barbara Dalmatiner, and Andreas Rohorzka for excellent technical support. We thank Nikolaus Osterrieder for generously providing the plasmid pEP-Kan-S for BAC mutagenesis.

Funding for this research was provided by a grant from the Austrian Science Fund (FWF) to I.G. (project number: P26420-B13). The funders had no role in study design, data collection and interpretation, or the decision to submit the work for publication.

## Additional Information

Supplementary Information is provided.

## Supplementary Table

**Supplementary Table 1.**
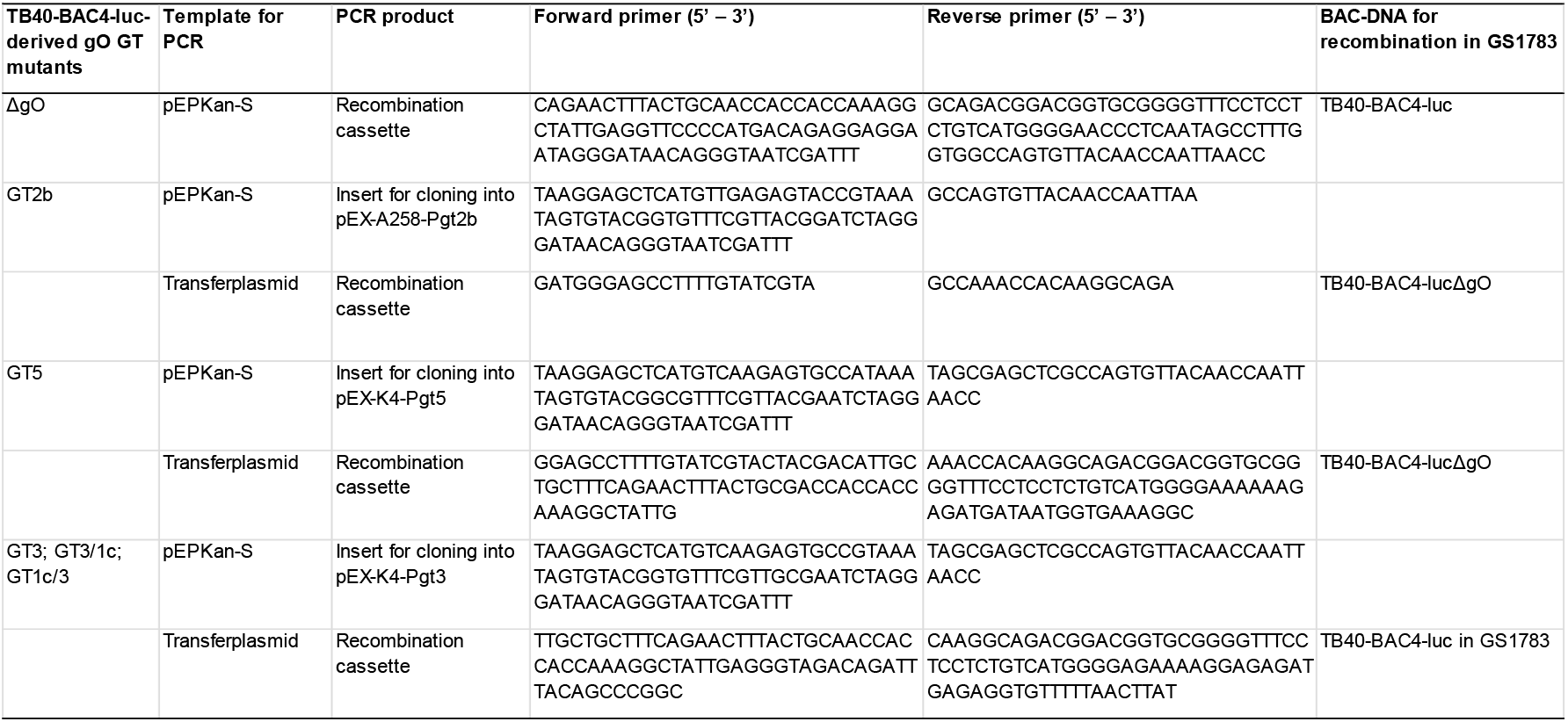
Primer sequences for en passant mutagenesis

## Supplementary figures

**Supplementary Figure S1:**
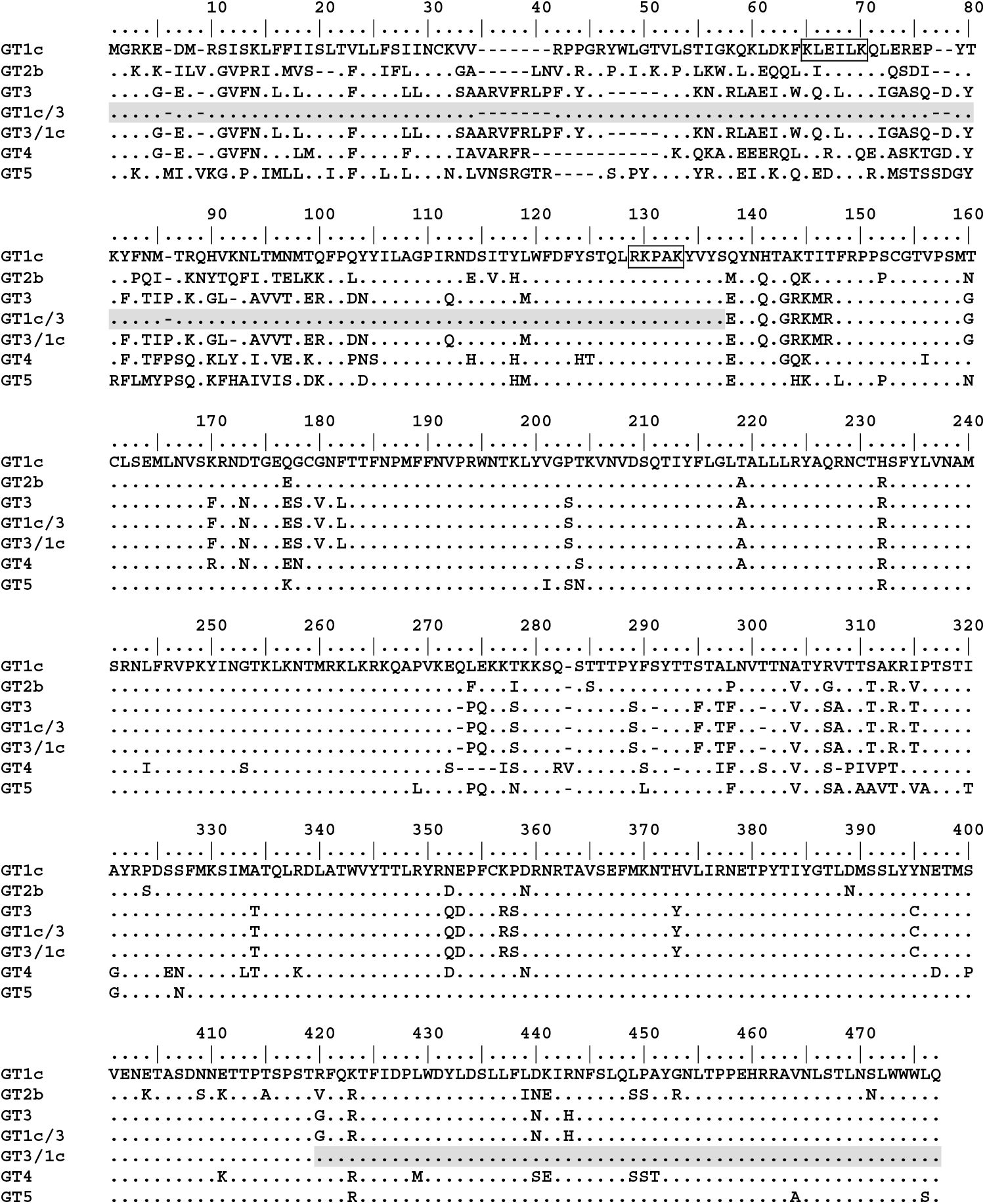
Amino acid alignment of gO genotype mutant sequences. Reference sequence of genotype (GT) 1c (TB40-BAC4; <u>ABV71596.1</u>) is aligned with GT2b (BE/29/2011; <u>AKI14139.1</u>), GT3 (HAN16; <u>AFR55727.1</u>), GT4 (Towne, <u>ACM48052.1</u>), gO GT5 (Merlin, <u>AAR31626.1</u>), and the two recombinant forms, GT1c/3 and GT3/1c. Putative PDGFRalpha binding sites as characterized recently (Stegmann et al., 2019) are depicted by black boxes. The grey-shaded regions of recombinant GT1c/3 and GT3/1c mutants indicate the GT1c sequence part.

**Supplementary Figure S2.**
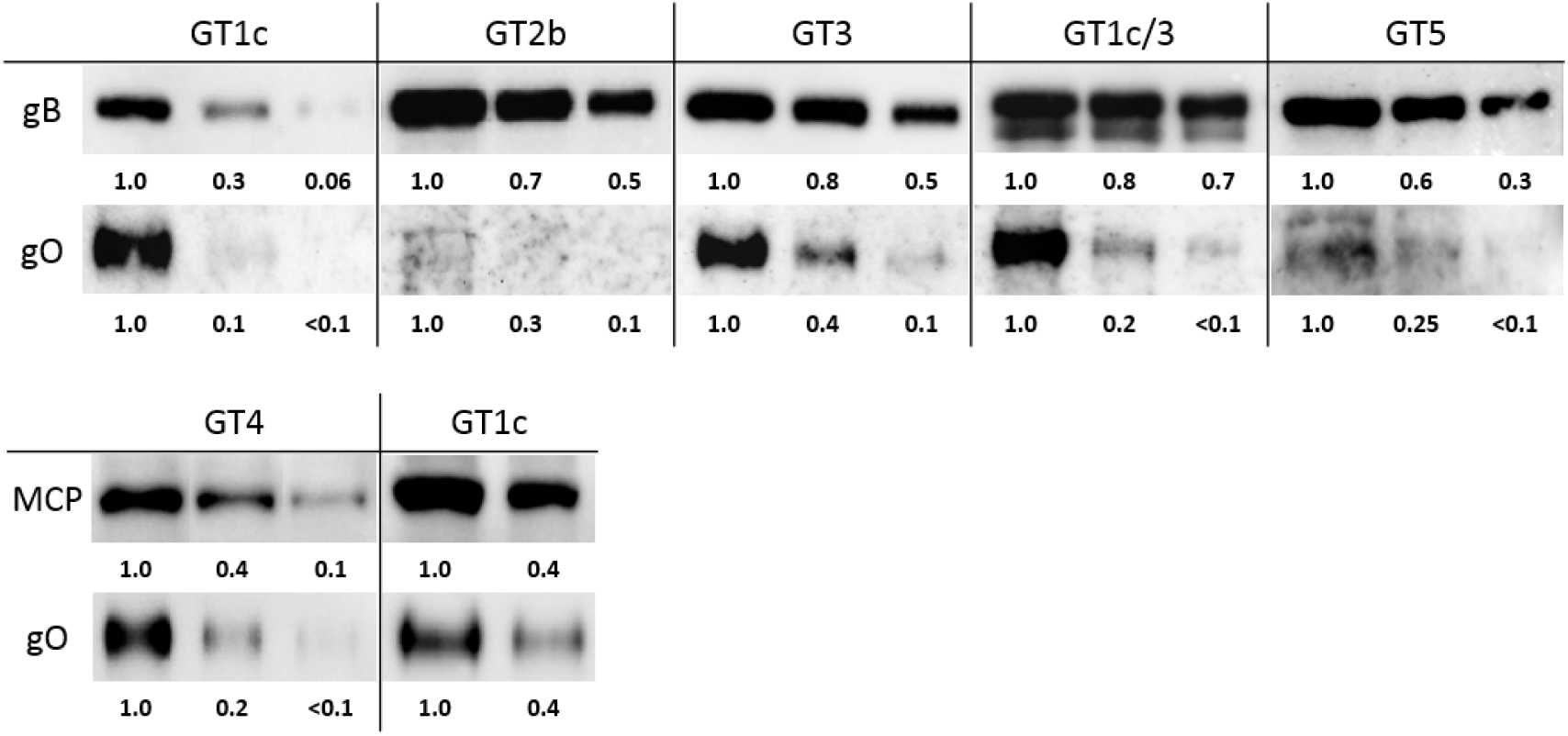
Reactivity of monoclonal anti-gO antibody, anti-gO.02 mAb, against distinct gO genotypic forms. Two to three 2-fold dilutions of virions were loaded on the gels. Reactivity against gO was compared to reactivity directed against gB (anti-gB mAb 2F12) or major capsid protein (MCP).

